# The anterior insular cortex in the rat exerts an inhibitory influence over the loss of control of heroin intake and subsequent propensity to relapse

**DOI:** 10.1101/2020.05.28.120725

**Authors:** Dhaval D. Joshi, Mickaël Puaud, Maxime Fouyssac, Aude Belin-Rauscent, Barry Everitt, David Belin

## Abstract

The anterior insular cortex (AIC) has been implicated in addictive behaviour, including the loss of control over drug intake, craving and the propensity to relapse. Evidence suggests that the influence of the AIC on drug-related behaviours is complex since in rats exposed to extended access to cocaine self-administration, the AIC was shown to exert a state-dependent, bidirectional influence on the development and expression of loss of control over drug intake, facilitating the latter but impairing the former. However, it is unclear whether this influence of the AIC is confined to stimulant drugs that have marked peripheral sympathomimetic and anxiogenic effects or whether it extends to other addictive drugs, such as opiates, that lack overt acute aversive peripheral effects. Thus, we investigated in outbred rats the effects of bilateral excitotoxic lesions of AIC, induced both prior to or after long-term exposure to extended access heroin self-administration, on the development and maintenance of escalated heroin intake and the subsequent vulnerability to relapse following abstinence. Compared to sham-surgeries, pre-exposure AIC lesions had no effect on the development of loss of control over heroin intake, but lesions made after a history of escalated heroin intake potentiated escalation and also enhanced responding at relapse. These data show that the AIC inhibits or limits the loss of control over heroin intake and propensity to relapse, in marked contrast to its influence on the loss of control over cocaine intake.

## Introduction

The anterior insular cortex (AIC) has been increasingly implicated in the pathophysiology, and associated behavioural manifestations of, drug addiction (Naqvi & Bechara, 2009; Verdejo-Garcia *et al.*, 2012). Chronic drug users have reduced AIC volume (Franklin *et al.*, 2002; Lyoo *et al.*, 2006; Ersche *et al.*, 2012; Ersche *et al.*, 2013; Lin *et al.*, 2019) and altered activity of the AIC-associated salience network (Goudriaan *et al.*, 2010; Wisner *et al.*, 2013; Clewett *et al.*, 2014; Sun *et al.*, 2017; Zilverstand *et al.*, 2018). Damage to the AIC in humans has been shown to result in a blunting of craving and withdrawal symptoms in individuals addicted to nicotine (Naqvi *et al.*, 2007; Abdolahi *et al.*, 2015).

In animal studies, selective bilateral excitotoxic lesions of the AIC curtail the escalation of cocaine self-administration (SA) in rats with a prolonged history of extended access to the drug and reduce cocaine-induced reinstatement of extinguished instrumental responses for the drug (Rotge *et al.*, 2017).

The behavioural consequences of damage to the insula in humans drug addiction have been suggested to reflect the role of this structure in interoception (Craig, 2003; Craig, 2009; Craig, 2011; Paulus & Stewart, 2014; Stewart *et al.*, 2019) and its associated influence on emotions (Craig, 2004; Wiens, 2005; Verdejo-Garcia *et al.*, 2006; Craig, 2010) and executive functions (Engstrom *et al.*, 2014), such as impulse control and decision-making (Weller *et al.*, 2009; Naqvi & Bechara, 2010; Jones *et al.*, 2011; Pattij *et al.*, 2014; Herman *et al.*, 2018; Rae *et al.*, 2019). Individuals cocaine (Vadhan *et al.*, 2009; van der Plas *et al.*, 2009; Cunha *et al.*, 2011), or heroin addiction (Rotheram-Fuller *et al.*, 2004; Lin *et al.*, 2012), as well as rats with a history of escalated cocaine SA, showed impairments in decision making tasks (Cocker *et al.*, 2019) that were similar to those following damage to the insula across species (Clark *et al.*, 2008; Daniel *et al.*, 2017). Similarly, a high impulsivity trait in humans (Lopez-Larson *et al.*, 2012) and rats (Belin-Rauscent *et al.*, 2015) is associated with decreased thickness of the AIC, and has also been shown across species to confer an increased tendency to escalate cocaine (Dalley *et al.*, 2007), but not heroin (McNamara *et al.*, 2010) intake and a vulnerability to switch from controlled to compulsive cocaine SA (Belin *et al.*, 2008; Ersche *et al.*, 2012).

In light of these studies, it remains to be established whether insula-dependent mechanisms, and associated interoceptive processes are involved in the loss of control over drug intake or whether their alteration by chronic drug exposure is an important factor influencing persistent drug use in addiction. In rats self-administering cocaine, the AIC has been shown to exert a bidirectional effect on the escalation of drug intake and its long-term maintenance. Bilateral excitotoxic lesions of the AIC in rats with well-established escalated drug intake, a measure of loss of control over intake (Ahmed & Koob, 1998; Ahmed *et al.*, 2000; DSM 5, American Psychiatric Association, 2013; Belin-Rauscent *et al.*, 2016), restored control over SA. However, similar lesions made prior to drug exposure markedly potentiated the escalation of cocaine intake (Rotge *et al.*, 2017).

Given the involvement of the insula in interoception, these differential effects of AIC lesions may reflect altered processing of the mixed appetitive and aversive effects of cocaine on initial exposure (Ettenberg & Geist, 1991; Ettenberg, 2004) and the change in the putative affective responses to cocaine over the course of the first two weeks of extended access to the drug (Klein *et al.*, 2020). Moreover, cocaine methiodide, a cocaine analogue that does not cross the blood-brain barrier, eventually acquires reinforcing properties in rats but only after a history of cocaine SA; it is not self-administered by drug naïve rats (Wise *et al.*, 2008; Wang *et al.*, 2013). Taken together, these data suggest that interoceptively-sensed peripheral consequences of self-administered cocaine mediated by the AIC may determine the loss of control over intake during long access to the drug and the subsequent maintenance of an escalated level of cocaine intake.

However, this role of the AIC may be restricted to stimulant drugs having distinctive peripheral and anxiogenic effects (Mitchell *et al.*, 1996). Opiates do not fully substitute for stimulant drugs in drug discrimination tasks (Negus *et al.*, 1998; Rowlett *et al.*, 2000; Rowlett *et al.*, 2004), suggesting differences in the peripheral versus central nature of the reinforcing effects of these different classes of drug.

Whether the AIC influences the loss of control over intake of opiates that lack the acute peripheral effects of stimulants after they have been self-administered has been investigated in the present study. The effects of bilateral excitotoxic lesions of the AIC on the establishment and maintenance of the escalation of heroin SA were studied in rats having 8-hours of daily extended access to the drug.

## Materials and methods

### Animals

48 male Sprague-Dawley rats (Charles River [Kent, UK]) weighing approximately 300g upon arrival were housed 4 per cage for a week of habituation to the vivarium. Rats had ad libitum access to food (standard chow) and water and were maintained under a reversed 12-hour light/dark cycle (light ON between 7.00pm and 7.00am). A week prior to SA, rats were food restricted to ~90% of their free-feeding body weight until the end of the experiment.

All experimental procedures were conducted under the project license 70/8072 held by David Belin in accordance with the United Kingdom Animals (Scientific Procedures) Act 1986, amendment regulations 2012 following ethical review by the University of Cambridge Animal Welfare and Ethical Review Body (AWERB).

### Study timeline

Rats were assigned to one of the two arms of this study, namely pre-exposure lesion and post-exposure lesion experiments, the timeline of which is illustrated in **Figure 1**. Each group underwent the same procedure to that used in our previous study with cocaine (Rotge *et al.*, 2017). For both experiments, after at least a week of habituation to the vivarium, rats were implanted with an indwelling catheter in the right jugular vein and, following a week, were trained to self-administer heroin over five 1-hour sessions under continuous reinforcement. They were then trained for 19 sessions under 8-hour extended access, conditions previously shown to promote robust escalation of SA (Ahmed *et al.*, 2000). Pre-exposure lesion rats underwent intracranial surgeries 5 days prior to the implantation of the intrajugular catheter while the post-exposure lesion rats underwent the surgical procedure after 12 days of training under extended access. These rats were also challenged for their propensity to relapse after a short 3-day period of abstinence. Following 3 days of forced abstinence after the 19^th^ extended access session, the post-exposure group underwent a relapse test where their tendency to extinguish instrumental responding for heroin and the reinstatement of the extinguished responding was tested. After finishing the relapse test, all rats from both experimental groups were sacrificed to harvest their brain for histological assessment of the AIC lesions.

**Figure 1:**
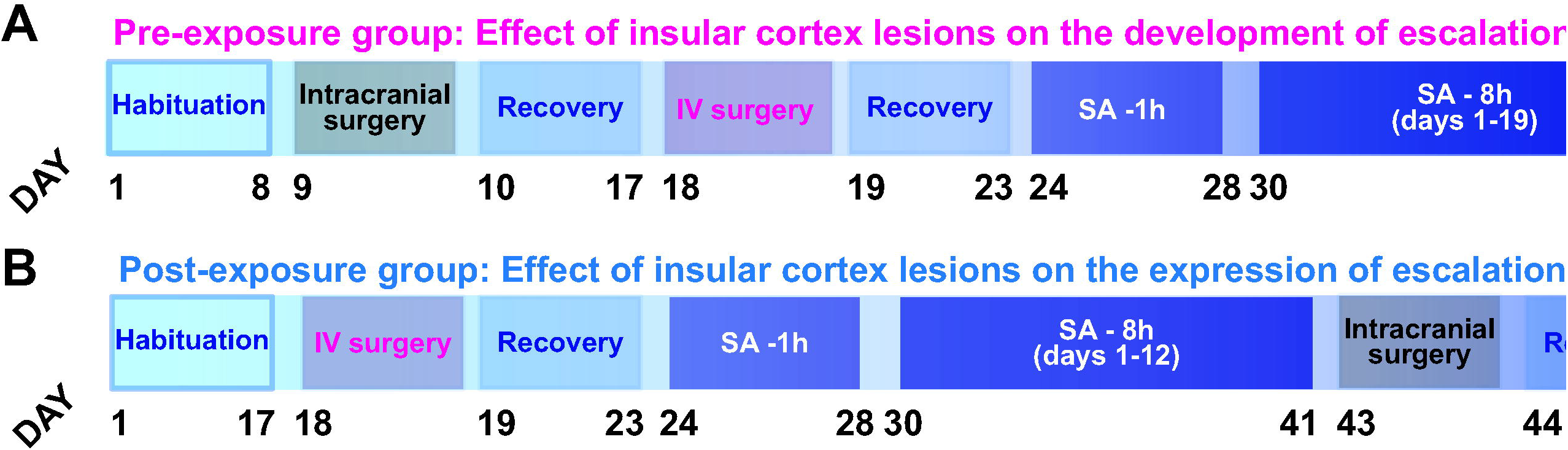
Timeline of the experiments. Two parallel studies were conducted to investigate the effects of bilateral lesions of the AIC on the establishment and maintenance of escalation of heroin SA. The general experimental design and timeline are similar to that used in our previous study with cocaine (Rotge *et al.*, 2017). All rats were first habituated to the vivarium for a week. **A)** For the pre-exposure lesion experiment, rats underwent sham or bilateral AIC lesion surgeries, and, following a period of recovery, were implanted with an indwelling catheter in the right jugular vein. At least five days later they were trained to acquire heroin SA under continuous reinforcement over five 1-hour daily sessions. They were then exposed to nineteen 8-hour extended access daily sessions prior to being sacrificed and their brains harvested for histological assessment of the lesions. B) for the post-exposure lesion experiment, rats were first implanted with an indwelling catheter in the right jugular vein, and after a period of at least 5 days of recovery, they were trained to acquired heroin SA under continuous reinforcement over five 1-hour daily sessions. Access was subsequently extended to 8 hours daily for 12 days, when rats escalated their heroin intake. These rats then received either sham or bilateral AIC lesions and, after a period of at least 8-9 days they were exposed to 7 additional extended access sessions. Following a three-day forced abstinence period rats were challenged in a 90-minute relapse test under extinction conditions. This was followed, as previously described (Belin *et al.*, 2009; Rotge *et al.*, 2017; Cocker *et al.*, 2019), by a 30‐minute non-contingent CS-induced reinstatement test, at the onset of which the heroin paired CS was presented noncontingently for 20 seconds. During the next 30‐minute period, active lever presses had no scheduled consequence. At the end of this 30‐minute period, three non-contingent infusions of increasing doses of heroin (20μg, 40μg and 80μg) were delivered at the beginning of the 30-minute periods of responding under extinction after which rats were sacrificed and their brains harvested for histological assessment of the lesions.

### Drugs

Heroin hydrochloride (MacFarlan-Smith, Edinburgh, UK) was dissolved in sterile 0.9% saline. Quinolinic acid (Sigma-Aldrich, Poole, UK) was dissolved in sterile PBS at a final concentration of 0.09 M.

### Stereotaxic surgery and bilateral AIC lesions

Stereotaxic surgeries were performed on a stereotaxic frame (WPI Hitchin, UK) under isoflurane anaesthesia (O_2_: 2 L/min; 5% for induction and 2-3% for maintenance) and analgesia (Metacam, 1 mg/kg, sc., Boehringer Ingelheim). The analgesic treatment was continued orally for three days post-surgery.

Quinolinic acid was infused bilaterally (0.8 μl/side) into the AIC at the following stereotaxic coordinates from the Paxinos and Watson atlas (2013): AP: +1.4, ML: ±5.2 from bregma, DV: −6.8 from skull. Infusions were performed as previously described (Rotge *et al.*, 2017) with 10 μl Hamilton syringes placed in a Harvard infusion pump and connected with a polyethylene tubing to 24 gauge injectors (Coopers needle works Ltd) at a rate of 0.35 μl/min. Injectors were left in place for 5 minutes following completion of the infusion to allow for diffusion away from the injector tip. Sham animals underwent the same procedure but with no infusion. All animals were allowed to recover for at least 10 days following surgery before testing resumed.

### Intra-jugular Surgery

Rats were kept under isoflurane anaesthesia (O_2_: 2 L/min; 5% for induction and 2-3% for maintenance and analgesia (Metacam, 1 mg/kg, sc., Boehringer Ingelheim) and implanted with a home-made indwelling catheter into their right jugular vein as previously described (Belin-Rauscent *et al.*, 2018). Following surgery, rats received daily oral treatment with the analgesic for three days and an antibiotic (Baytril, 10mg/kg, Bayer), which they first received on the day prior to surgery, for a week. Catheters were flushed with 0.1 ml of heparinized saline (50 U/ml, Wockhardt®) in sterile 0.9% NaCl every other day after surgery and then before and after each daily SA session. Following this surgery all rats were single housed until the end of the experiment (as illustrated in **Figure 1**).

### Self-administration training

SA experiments were conducted in 24 standard operant chambers (Med Associates Inc., St. Albans, VT, USA) controlled by Whisker software (Cardinal & Aitken, 2010). Chiefly each (29.5 × 32.5 × 23.5 cm) chamber was housed in ventilated, sound-attenuating cubicles. Sidewalls were aluminium; the ceiling, front and back walls were clear polycarbonate. Two retractable levers (4 cm wide) were surmounted 8cm above the grid floor and 12 cm apart, with a white cue light (2.5W, 24V) and a white house light (2.5W, 24V), situated on the wall opposite the levers. Implanted catheters were connected to a 10 ml syringe driven by an infusion pump (Semat Technical, Herts, UK) via Tygon tubing itself protected within a spring leash attached to a swivel connected to a balanced metal arm secured outside of the chamber.

Each animal was permanently assigned an active lever (either left or right) and this was counterbalanced across animals. During the SA sessions, each trial began with the house light being turned on and the two levers being inserted.

Active lever presses were reinforced under a fixed ratio 1 schedule of reinforcement by the delivery of a bolus of heroin hydrochloride (40μg/100μl/1.7s) concomitant to a 20-second time-out period where the levers were retracted, the house lights were turned off and a light was presented over the active lever as a conditioned stimulus (CS). Meanwhile, inactive lever presses had no programmed consequences.

Rats acquired heroin SA over five 1-hour short access sessions and were subsequently exposed to 8-hour long extended access sessions until the end of training.

Three rats were excluded from the analyses, one because it failed to complete the SA training due to catheter failure, two others because they were sacrificed before the end of the training following autotomy.

### Relapse test

The relapse test was designed as previously described (Belin-Rauscent *et al.*, 2018). After 3 days of abstinence, rats were given the opportunity instrumentally to respond for heroin for 90 minutes in the context they had previously been trained to self-administer the drug, but under extinction conditions (i.e. no heroin was infused) (Belin-Rauscent *et al.*, 2018) (**Figure 3D**). This was followed, as previously described (Belin *et al.*, 2009; Rotge *et al.*, 2017; Cocker *et al.*, 2019), by a 30‐minute non-contingent CS-induced reinstatement test, at the onset of which the heroin paired CS was presented noncontingently for 20 seconds. During the next 30‐minute period, active lever presses had no scheduled consequence. At the end of this 30‐minute period, three non-contingent infusions of increasing doses of heroin (20μg, 40μg and 80μg) were delivered at the beginning of 30-minute periods of responding under extinction.

### Histology and lesion assessment

Animals were anaesthetised with pentobarbital (Euthatal, Merial, 750 mg/Kg) and transcardially perfused with 0.01M phosphate-buffered saline (PBS), followed by 4% neutral-buffered formaldehyde (NBF). Brains were harvested and post-fixed for at least 24h at 4°C in 4% NBF. Brains were then cryo-protected in a 30% sucrose solution (prepared in PBS 0.01M). After a quick freezing on dry-ice, brains were processed with a cryostat (Leica Microsystems) into 50μm coronal sections that were placed directly on gelatine-coated slides. Brain slices were then stained with cresyl-violet to allow for visualisation of lesions and covered with cover slips using a DPX mounting medium. Lesion assessment was performed blind to the behavioural data by two independent experimenters from images taken with a bright-field microscope (Zeiss Axio Imager Z2, Zeiss, Oberkochen, Germany) illustrated in **Figure 2C**.

**Figure 2:**
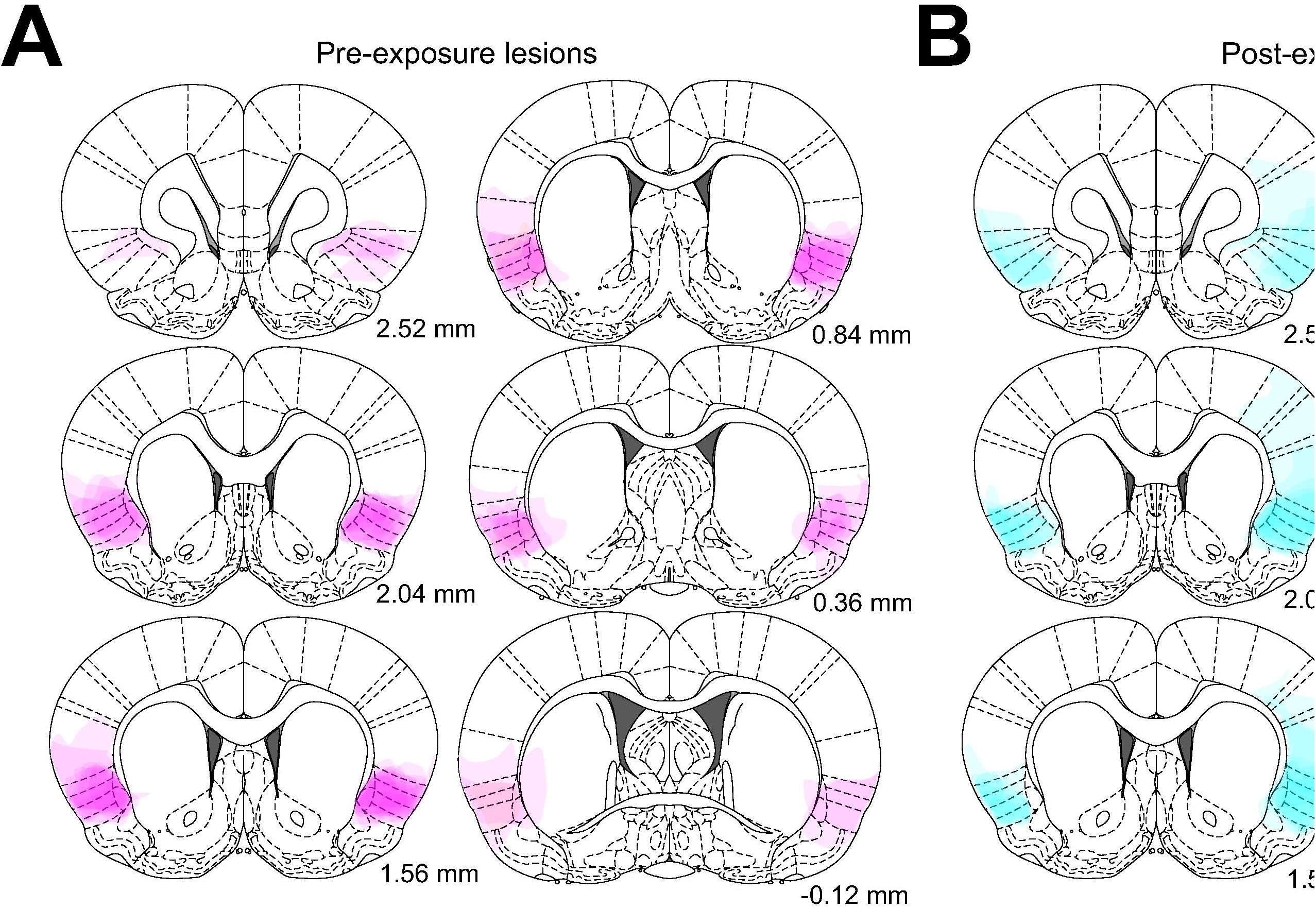
Bilateral lesions encompassed the entirety of the AIC. Lesion profiles of the pre-(**A**) and post-exposure (**B**) groups illustrated. The co-ordinates relative to bregma are noted under each brain atlas template (extracted from Paxinos & Watson, 2013). **C**. Exemplar bright-field image of a Cresyl Violet-stained, 50 μm-thick coronal section imaged at 10X magnification with an outline of the lesion drawn in red.

Five rats were excluded from analyses because the lesions spread significantly outside the AIC across the length of the anterior-posterior plane.

### Data and statistical analyses

All statistical analyses were performed using Statistica 10 (StatSoft, Inc, 2011). Data are presented as means ± SEM or box plots [medians ±25% (percentiles) and min/max as whiskers)].

Assumptions for normality of distribution and homogeneity of variance were verified using the Kolmogorov-Smirnov and the Cochran C test, respectively. Where normality and/or homogeneity of variance was substantially violated, data were log transformed.

A two-sample t-test was used to compare total active lever presses during the relapse test across groups of rats and results are displayed as t-statistic (degrees of freedom) and two-tailed p-value.

A repeated measures analysis of variance (RM-ANOVA) was used to analyse heroin infusions across days and active lever presses across time bins of the relapse test with SA days or relapse test time bins as within-subject factors and treatment (“lesioned” vs “sham”) as between-subject factor.

The analysis of the influence of AIC lesions on the expression of escalated heroin SA in the post-exposure group was carried out using a RM-ANOVA with a block design in which the was last 6 days before and after the lesion were used as pre and post lesion blocks (**Figure 3C**). The potential reliance of performance at relapse, under extinction conditions, on previous between-group differences in responding under reinforcement was measured using regression with total active lever presses before lesions and total active lever presses after lesions as predictor variables and the total active lever presses in the first 5 minutes of relapse as the dependent variable. The total lever presses emitted during the first 5 minutes were chosen as we wanted to investigate the differences in responding during relapse at a point they were the most pronounced.

**Figure 3:**
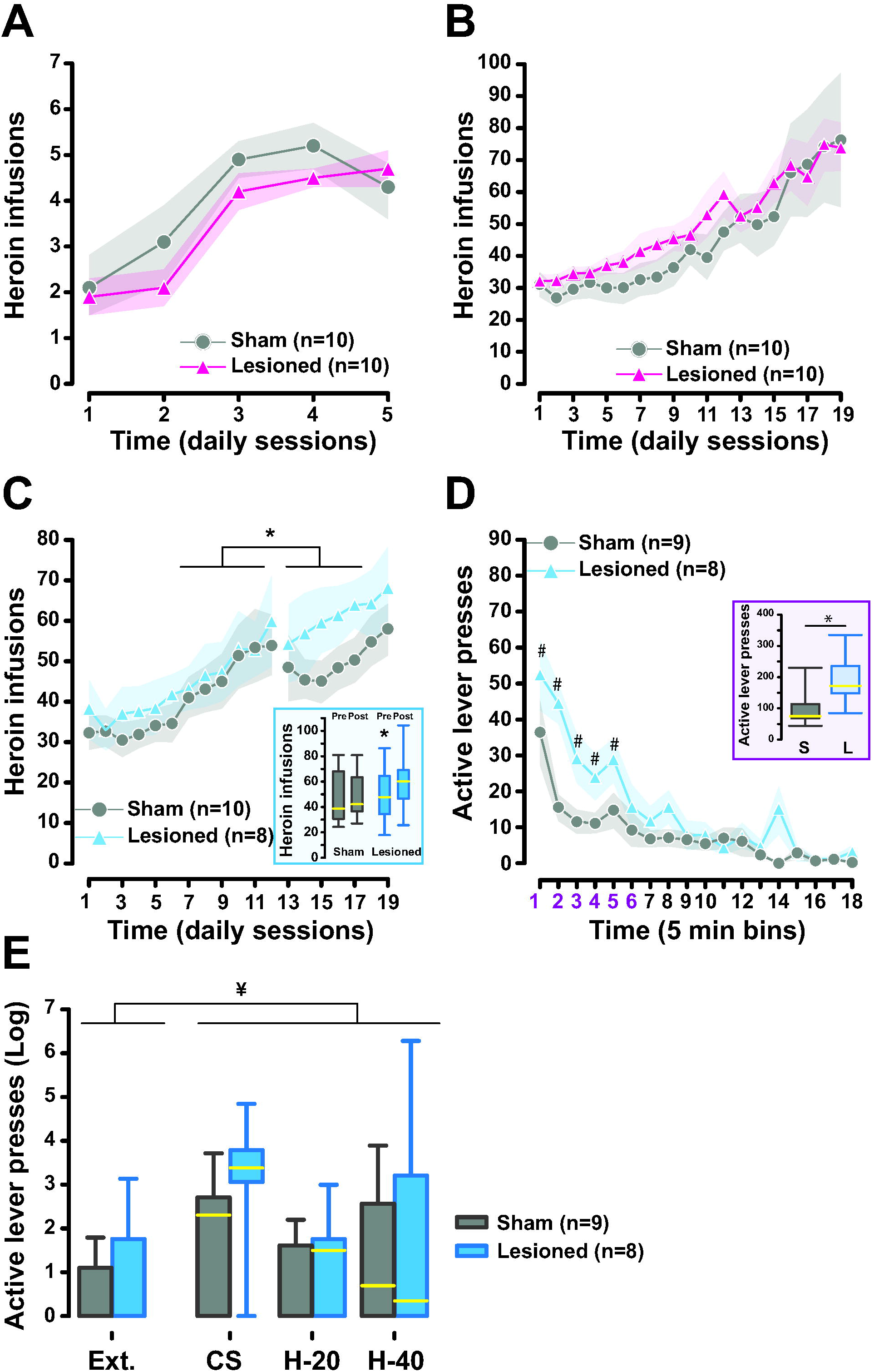
Post-heroin exposure AIC lesions potentiate the expression of escalated heroin intake and subsequent propensity to relapse. Pre-exposure lesions of the AIC had no effect on the acquisition of heroin SA (**A**). Both sham and AIC lesioned rats acquired heroin SA under continuous reinforcement at similar rates over 5, 1-hour daily sessions. Similarly, pre-exposure AIC lesions had no effect on the escalation of heroin SA over the course of 19, 8-hour daily extended access sessions (**B**). In marked contrast, post-escalation AIC lesions resulted in further escalation of heroin intake compared to the control group (**C**). Thus, while sham rats re-attained pre-surgery levels of intake, AIC lesioned rats furthered their increased their intake eventually to reach levels that were higher than that those attained prior to the lesion [*: main effect of group x block interaction: F_1,16_ = 5.39, p = 0.03, pη^2^ = 0.25]. The break in the data trace represents the time of lesion surgery. The inset displays the average No of infusions over 6-day blocks before and after surgeries in lesioned and sham animals. Bilateral AIC lesions also potentiated the individual tendency to relapse after short-term abstinence (**D**). AIC-lesioned rats showed increased responding during a 90-minute relapse challenge under extinction [main effect of time bin x lesion group interaction F_17,255_ = 2.20, p = 0.005, pη^2^ = 0.13], which was observed mainly during the first 30 minutes of the challenge test (insert: [* indicates t_15_ = 2.78, p = 0.019, two-sample t-test (2-tailed)]. Thus, while both groups reached extinguished levels of responding by the last 15 minutes of the test, the instrumental performance of sham animals only differed from extinguished levels during the first five minutes of the test, whereas responding by AIC lesioned rats remained significantly higher for 20 minutes longer (#: different from the last 15 min, Newman-Keuls post-hoc). This increased responding during the relapse challenge under extinction induced by AIC lesions was not seen during CS- or heroin-induced reinstatement challenges. While the non-contingent CS and heroin presentations equally increased responding as compared to baseline (¥: reinstatement blocks vs baseline: F_1,15_ = 10.31, p < 0.01], no differences were between AIC lesioned and sham rats were observed.

For the reinstatement test, in a similar manner to previously described (Rotge *et al.*, 2017; Cocker *et al.*, 2019) instrumental performance in response to CS or drug presentation decreased throughout each 30‐minute block, such that animals had extinguished responding towards the end of each block, a point at which their performance does not differ from that they had reached at the end of the preceding 90-minute extinction period except for the second heroin-primed reinstatement block during which rats did not extinguish at all [main effect of block: F_3,45_ = 2.8153, p = 0.049], thereby preventing the assessment of the influence of the third infusion on responding. Consequently, we discarded that third heroin-primed reinstatement block from the analysis. Thus, the reinstatement of instrumental responding primed by non-contingent presentation of either a CS or non-contingent infusions of heroin was measured over the first 10 minutes of each 30-minute block in comparison to the last 10 minutes of extinction considered here as baseline. This strategy enabled us to measure the influence of non-contingent CS or drug presentations on the actual reinstatement of an extinguished response without the confounding influence of the extinction process that takes places over each 30-minute block [main effect of block: F_2,30_=2.45, p=0.1 and block x lesion interaction: F_2,30_<1 for the performance over the last 10 minutes of each block].

Upon significant main effects, differences between individual means were analysed using the Newman-Keuls post-hoc test. For all analyses, statistical significance was accepted at α = 0.05. Effect sizes are reported as the partial η^2^ (pη^2^).

## Results

Excitotoxic lesions targeting the AIC were overall restricted to the insula, as shown in **Figure 2**. In three individuals of the post-exposure lesion group, the lesion was observed to have spread dorsally to the somatosensory cortex, but only unilaterally and on a restricted portion of the antero-posterior plane. Since the lesion spread was only unilateral and these rats did not differ behaviourally from the other animals of this group they were included in the subsequent analyses.

Bilateral AIC lesions did not influence the propensity to acquire heroin SA under continuous reinforcement in that rats from the sham and pre-exposure lesion groups did not differ in the progressive increase in their heroin infusions over the five 1-hour daily sessions [main effect of session: F_4, 72_ = 19.29, p < 0.001, pη^2^ = 0.52, lesion: F_1, 18_ = 0.61, p = 0.44, pη^2^ = 0.03 and lesion x session interaction: F_4, 72_ = 0.86, p = 0.49, pη^2^ = 0.05] (**Figure 3A**). Upon the subsequent introduction of 8-hour long-access sessions, both AIC lesioned and sham rats displayed a robust escalation of intake from initially 30 to 70 infusions per session within ~20 days [main effect of session: F_18, 324_ = 14.77, p < 0.001, pη^2^ = 0.45, lesion: F_1, 18_ = 0.35, p = 0.56, pη^2^ = 0.02 and lesion x session interaction: F_18, 324_ = 0.38, p = 0.99, pη^2^ = 0.02] (**Figure 3B**). Thus, pre-exposure AIC lesions did not influence the establishment of loss of control over heroin SA.

In contrast, post-exposure bilateral AIC lesions performed once rats had developed escalation over 12 days of exposure to extended access to the drug [main effect of session: F_11, 176_ = 9.32, p < 0.001, pη^2^ = 0.37 and session x lesion interaction: F_11, 176_ = 0.22, p = 0.99, pη = 0.01] resulted in a prolonged potentiation of subsequent escalated intake of heroin [main effect of block: F_1, 16_ = 7.49, p = 0.01, pη^2^ = 0.32, lesion: F_1, 16_ = 0.47, p = 0.50, pη^2^ = 0.03 and block x lesion interaction: F_1, 16_ = 5.39, p = 0.03, pη^2^ = 0.25] (**Figure 3C**). Thus, while sham rats showed an escalated level of intake post-surgery that was similar to that reached pre-surgery, thereby establishing a plateau characteristically reached by day 12 of extended access (Ahmed *et al.*, 2000; McNamara *et al.*, 2010), lesioned rats instead showed a continuing escalation of intake eventually reaching higher levels of intake than that established prior to the lesion (**Figure 3C** inset).

This differential influence of AIC lesions on heroin intake was not due to an effect on body weight [main effect of block × lesion interaction: F_1,16_ = 0.94, p = 0.35, pη^2^ = 0.06]. However, the exacerbation of heroin escalation by bilateral AIC lesions was related to an increased tendency subsequently to persist in responding for the drug at relapse, when rats where challenged to respond for 90 minutes under extinction in the SA context following a short period of forced abstinence [main effect of time: F_17, 255_ = 16.42, p < 0.001, pη^2^ = 0.52 and time x lesion interaction: F_17, 255_ = 2.20, p = 0.005, pη^2^ = 0.13] (**Figure 3D**). Thus, although AIC lesioned rats eventually reached levels of responding similar to that of sham animals by the end of the 90-minute relapse challenge session under extinction, they showed much higher levels of responding during the first 30 minutes of the relapse challenge [t_15_ = 2.78, p = 0.019, two-sample t-test (2-tailed)] (**Figure 3D** and **Figure 3D** inset). This differential engagement in responding at relapse was further shown by the observation that initial performance in sham animals no longer differed from that of its extinguished level, measured over the last 15 minutes of the challenge, after 5 minutes whereas responding by AIC lesioned remained different from extinguished levels for 25 minutes.

The increased responding at relapse shown by AIC lesioned rats was not attributable to their higher instrumental output under reinforcement following surgery (**Figure 3C**). Indeed, performance during the initial phase of the relapse challenge was not better predicted by the total level of lever pressing during the post-lesion period (adjusted R^2^ = 0.39, p = 0.004) than by that during the pre-lesion period (adjusted R^2^ = 0.32, p = 0.01). In contrast with their influence on the motivation to respond at relapse, AIC lesions had no effect on CS- or heroin-induced reinstatement (**Figure 3E**). Thus, while non-contingent presentation of the CS or heroin induced the anticipated increase in instrumental responding in all rats as compared to baseline performance over the last 10 minutes of the 90-minute extinction session [planned comparison: F_1,15_ = 10.31, p < 0.01], this reinstatement of extinguished instrumental responding was not quantitatively different between AIC lesioned and sham rats [main effect of session x lesion interaction: F_1,15_ = 3.62, p = 0.08, pη^2^ = 0.19 and F_2,30_ = 0.05, p = 0.95, pη^2^ = 0.003 for CS- and heroin-primed blocks, respectively] (**Figure 3E**).

## Discussion

The results of the present study show that pre-exposure bilateral excitotoxic lesions of the AIC have no effect on the acquisition of heroin SA or the establishment of the escalation of heroin intake that develops over 19 days of extended access. In marked contrast, rats with post-training AIC lesions, made once they have reached a plateau of escalated intake (Ahmed *et al.*, 2000; McNamara *et al.*, 2010) that usually remains stable for weeks (McNamara *et al.*, 2010), continued to escalate their intake whereas sham-operated control rats maintained the pre-lesion plateau. The enhanced escalation following AIC lesions was accompanied by an increased propensity to relapse, as measured as a higher rate of responding for the drug under extinction following a brief period of abstinence (Belin-Rauscent *et al.*, 2018).

The effects of post-escalation AIC lesions on the further escalation of heroin intake could not be attributable to an effect on body weight, which influences the pharmacokinetic properties of the drug (Rook *et al.*, 2006a; Rook *et al.*, 2006b), or an increased tolerance to the reinforcing properties of heroin since AIC lesioned rats did not differ from sham controls in their increase in body weight following AIC lesions, or in their response to heroin-primed reinstatement of responding after abstinence. This is further suggested by the absence of any increase in intake in rats receiving AIC lesion prior to either brief or protracted extended access heroin SA sessions. The increased responding at relapse shown by AIC lesioned rats was not specifically predicted by their higher level of responding under reinforcement over the 7 sessions of post-lesion SA. Nor was it predicted by the behavioural manifestation of impaired extinction learning (Peters *et al.*, 2009; Millan *et al.*, 2011) since they reached similar levels of responding by the end of the 90-minute session and each of the 30-minute reinstatement blocks as did sham controls.

The post-escalation, unlike the pre-escalation, AIC lesioned rats, experienced a prolonged period of abstinence that necessarily followed surgery before resuming heroin SA. However, had this period of abstinence alone facilitated increased intake when rats resumed self-administering heroin, as might have been expected, this would have been seen in both lesioned and control rats. However, only AIC lesioned rats further escalated their intake, while controls maintained their pre-operatively acquired escalated intake plateau.

Taken together, these findings suggest that the AIC is not directly involved in the neural mechanisms underlying the escalation of heroin intake, previously shown to involve the endogenous stress-system (Greenwell *et al.*, 2009; Schlosburg *et al.*, 2013; Park *et al.*, 2015), but that it instead is involved in the establishment of an allostatic level of intake in rats during long access SA sessions (Koob & Moal, 1997) and without which rats continue to escalate their intake.

The inhibitory influence the AIC exerts over escalated heroin intake is accompanied by a similar influence on the propensity to relapse, as measured by responding under extinction conditions after a short period of abstinence, but not on the incentive properties of the drug or its associated CS as measured as drug- or CS-induced reinstatement of an extinguished instrumental response (Belin *et al.*, 2009). These results are in contrast to our previous demonstration of bidirectional control by the AIC over the development and maintenance of the escalation of cocaine SA, whereby pre-exposure AIC lesions enhanced the initial escalation of cocaine intake, while post-escalation AIC lesions restored control over cocaine intake (Rotge *et al.*, 2017).

The contrasting effects on heroin and cocaine SA may reflect that the interoceptive function attributed to the AIC is differentially engaged in the establishment and maintenance of loss of control over stimulants, that have marked peripheral sympathomimetic effects, and opiates that do not. This is evidenced by the finding that stimulant drugs and opiates do not necessarily substitute for each other in drug discrimination tasks (Dykstra *et al.*, 1992) and, when they do, they do so asymmetrically. Hence, opiates can substitute for the μ-opiate receptor-dependent component of the discriminative properties of cocaine (Rowlett *et al.*, 2000) and can potentiate the discriminative properties of cocaine (Suzuki *et al.*, 1995). Cocaine, by contrast, does not substitute for, and does not influence, the discriminative properties of opiates. Furthermore, tolerance to the discriminative properties of stimulants is dissociable from those of tolerance to morphine (Young *et al.*, 1992).

Cocaine has marked anxiogenic discriminative effects (Shearman & Lal, 1981) and triggers anxiogenic states (Raven *et al.*, 2000; Ettenberg, 2004; Guzman & Ettenberg, 2007; Schank *et al.*, 2008) by increasing blood pressure and inducing tachycardia. The demonstration that cocaine methiodide, a peripherally active but centrally inactive, cocaine analogue is reinforcing and self-administered only in rats having previously acquired cocaine SA (Wang *et al.*, 2013) suggests that rats must learn to associate the aversive peripheral effects of cocaine with its central actions, that together constitute cocaine’s reinforcing properties (Everitt & Robbins, 2005), for the peripherally-generated cocaine-conditioned stimuli to be associated with cocaine’s central reinforcing properties.

The different effects of AIC lesions on the escalation of cocaine versus heroin intake likely reflect the differential engagement of the AIC in processing the powerful, anxiogenic sympathomimetic peripheral effects of cocaine, but its lack of engagement by the peripheral effects of heroin, which is CNS-depressant but not aversive. Hence, the influence of the AIC and the interoceptive mechanisms it subserves on heroin SA, escalation and relapse may be of a very different nature to its influence on cocaine reinforcement mechanisms. This is further indicated by the effects of AIC manipulations on stimulant (Contreras *et al.*, 2012), but not opiate (Li *et al.*, 2013), conditioned placed preference.

The impaired ability of post-escalation AIC lesioned rats to reach a stable higher plateau of heroin intake when given extended access may reflect disrupted drug satiety (Steidl *et al.*, 2015) in an allostatic state in which intake is regulated and maintained around an elevated baseline (Koob & Moal, 1997). The reinforcing and motivational properties of opiates have been shown to depend on systemically released peptides involved in food satiety (Douton *et al.*, 2019) and acute food restriction-induced heroin seeking depends on leptin and ghrelin (Shalev *et al.*, 2001; D’Cunha *et al.*, 2020). Chronic exposure to opiates triggers long-term adaptations in satiety-controlling (Duraffourd *et al.*, 2012), enteric neurons (Duraffourd *et al.*, 2014) that project via visceral afferent pathways to the insula (Mayer, 2011). On this background, we propose that the AIC may play an analogous role in regulating allostatic opiate intake to maintain a balance between the incentive and satiety effects of heroin around the higher set-point that is established by escalation during extended access to the drug.

The contrasting consequences of damage to the AIC on the development and maintenance of opiate and stimulant escalated intake is further evidence that the mechanisms of opiate and stimulant addiction are more dissimilar than initially proposed (Badiani *et al.*, 2011).

## Acknowledgements

This work, carried out at the Department of Psychology, University of Cambridge, was funded by a Programme Grant from the Medical Research Council to BJE and DB (MR/N02530X/1) and a research grant from the Leverhulme Trust to DB (RPG‐2016‐117). DJ is supported by a BBSRC-DTP/Shionogi joint grant to DB (G101457).

DB, MP, and MF designed the experiment. MP and MF carried out all the behavioural procedures. DB performed the IV surgeries and MP performed the IC surgeries. MP, ABR and DDJ performed post mortem assessments of the lesions and data analysis. DDJ and DB performed the statistical analyses and wrote the manuscript together with BJE.

Heroin hydrochloride was provided to DB by the NIDA Drug Supply Programme. The authors would like to thank Aïcha Aleian for her help with half the behavioural experiments.

